# Axonal mRNA in human embryonic stem cell derived neurons

**DOI:** 10.1101/066142

**Authors:** Rebecca L. Bigler, Joyce W. Kamande, Raluca Dumitru, Mark Niedringhaus, Anne Marion Taylor

## Abstract

The identification of axonal mRNAs in model organisms has led to the discovery of multiple proteins synthesized within axons for functional roles such as axon guidance and injury response. The extent to which protein synthesis within the axon is conserved in humans is unknown. Here we used axon-isolating microfluidic chambers to characterize the axonal transcriptome of human embryonic stem cells (hESC-neurons) differentiated using a protocol for glutamatergic neurons. Using gene expression analysis, we identified mRNAs proportionally enriched in axons, representing a functionally unique local transcriptome as compared to the human neuronal transcriptome inclusive of somata and dendrites. Further, we found that the most abundant mRNAs within hESC-neuron axons were functionally similar to the axonal transcriptome of rat cortical neurons. We confirmed the presence of two well characterized axonal mRNAs in model organisms, β-actin and GAP43, within hESC-neuron axons using multiplexed single molecule RNA-FISH. Additionally, we report the novel finding that oxytocin mRNA localized to these human axons and confirmed its localization using RNA-FISH. This new evaluation of mRNA within human axons provides an important resource for studying local mRNA translation and has the potential to reveal both conserved and unique axonal mechanisms across species and neuronal types.

## INTRODUCTION

Intra-axonal translation is involved in multiple cellular functions, including axon maintenance, guidance, synaptogenesis, presynaptic plasticity and response to injury^1-5^. Specific axonal mRNAs implicated in these neuronal functions have been experimentally identified in *Aplysia* sensory neurites, rodent peripheral and central neurons, and multiple other model systems^2, 6-10^. In axons and synapses, which are remote from the cell body, local translation may be critical for spatially restricted and rapid responses to extracellular signals such as glutamate, brain derived neurotrophic factor, neurotrophin-3 and netrin-1^8, 11-14^.

Comparing axonal transcriptomes of various neuronal subtypes has the potential to highlight conserved roles of intra-axonal translation and reveal unique functions and targets within each neuron type. Direct evaluation of primary human neurons is severely restricted due to the limited ability to obtain tissue or culture isolated neurons. Human stem cell derived neurons are a biologically relevant and tractable *in vitro* system and protocols have been developed to differentiate stem cells into specific neuronal subtypes, such as motor, striatal, dopaminergic and glutamatergic neurons ^15-20^. Here we sought to identify the axonal transcriptome of human embryonic stem cells differentiated into neurons enriched for glutamatergic lineage^20, 21^ and compare this transcriptome to axonal transcriptomes from rodent model systems.

## RESULTS

### Differentiation of human embryonic stem cell derived neurons in microfluidic chambers

To identify transcripts within human axons we first modified an existing microfluidic chamber design for compartmentalizing and harvesting pure axons, originally developed for murine neurons^22^. To optimize the growth of hESC-neurons, which have an increased nutrient demand compared to primary rodent neurons, we increased the height of the fluidically isolated compartments to provide a greater volume of media surrounding the cells. In these microfluidic chambers, neurons were seeded in the somatic compartment and stochastic process growth resulted in very long and fine processes passing through the microgrooves to the axonal compartment (Fig. 1A). Similar chamber designs have been used to characterize the axonal transcriptome of primary rat cortical neurons and mouse motoneurons and for biochemical analysis of pure axon^23, 24^.

**Figure 1.**
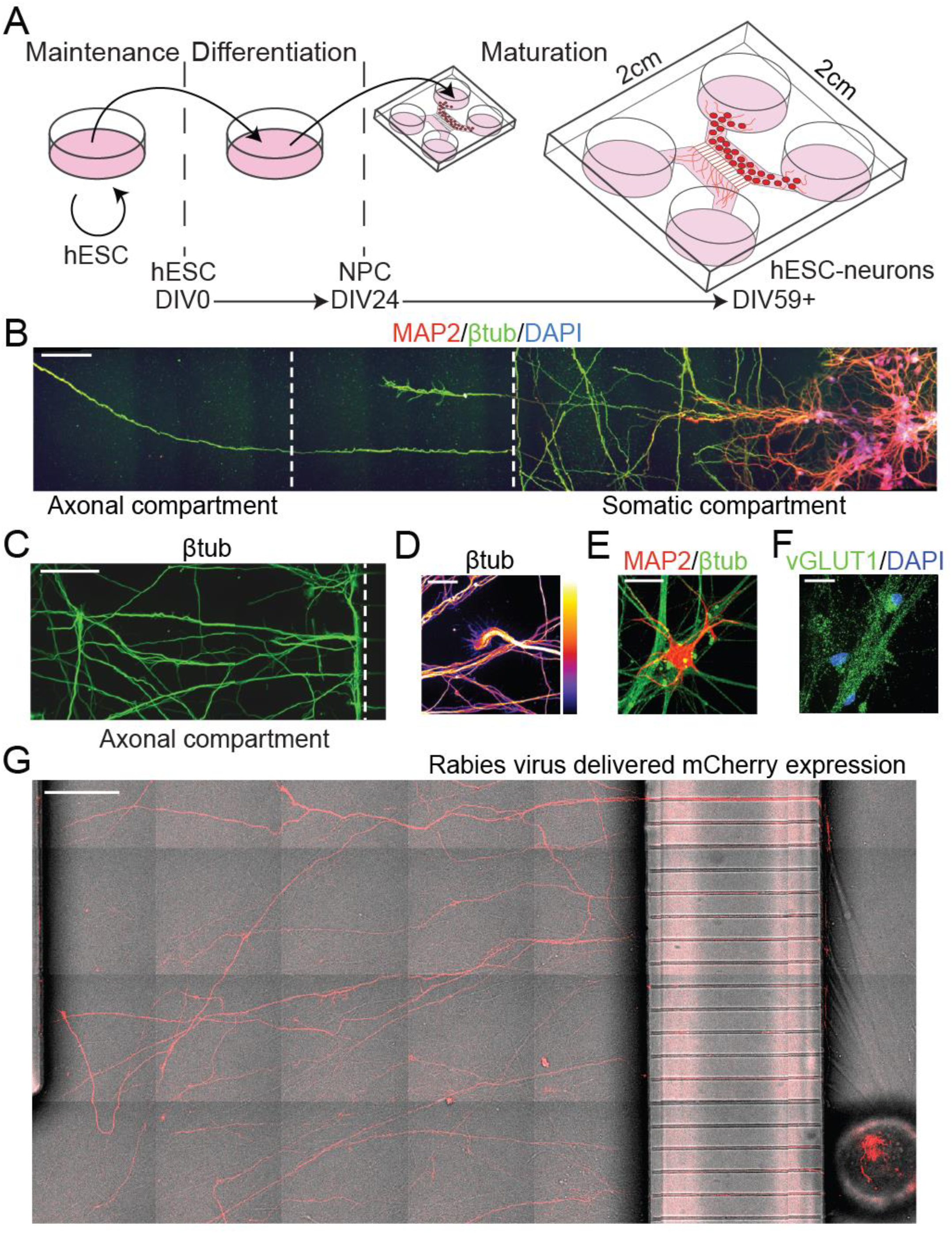
Human embryonic stem cell derived neurons (hESC-neurons) matured in axon isolating microfluidic chambers. (A) Schematic and timeline of hESC-neuron differentiation and maturation in microfluidic chambers. (B) Montage image of MAP2 (dendrite), β-tubulin III (βtub, axon) and DAPI immunostaining spanning the somatic and axon compartments of DIV39 hESC-neurons cultured within a microfluidic chamber. White dashed lines delineate the boundaries of the microgroove barrier. (C) Representative montage image of β-tubulin III immunostaining of axons in the axon compartment on DIV58. (D) Image of a β-tubulin III immunostained hESC-neuron growth cone within the axon compartment, “Fire” LUT pseudocolor pixel intensity of 0 is black and of 255 is white. (E) hESC-neurons exhibited dendritic arborization revealed by MAP2 immunostaining. (F) Matured hESC-neurons were positive for the glutamatergic marker VGLUT1. (G) A modified rabies virus encoding the mCherry fluorescent protein was applied to the axonal compartment for 2 h to infect neurons with axons extending into the axonal compartment and imaged 2 days after infection (DIV63). Scale bars are: 100 μm (B, C), 25 μm (D, E, F) and 200 μm (G).

Differentiation of hESC-neurons began in traditional tissue culture dishes on days *in vitro* (DIV) 0 (Fig. 1A) according to an established protocol to generate functionally matured glutamatergic neurons that produce action potentials after approximately 6 weeks of differentiation^20, 21^. On DIV24 neural progenitor cells were transferred to our custom axon isolating microfluidic chambers and matured to hESC-neurons over the next 30 days. As early as DIV39 long, neurites could be seen in the microgrooves; Microtubule-associated protein 2 (MAP2) and Neuron-specific class III β-tubulin (β-tubulin III, βtub) staining, markers for cell bodies and axons respectively, suggest that the majority of neurites entering the axonal compartment are axons (Fig. 1B). β-tubulin III staining revealed extensive neurite growth within the axonal compartment at DIV58 (Fig. 1C) as well as healthy growth cones (Fig. 1D). After five weeks in culture, some clustering of somata occurred, as is common in long term cultures of hESC-neurons. Cells demonstrated neuron arborization as revealed by MAP2 staining (Fig. 1E). After DIV49 >90% of cells within the microfluidic chamber differentiated into neurons, 198 of 209 of DAPI-labeled nuclei were positive for MAP2 and/ or β-tubulin III. The remaining cells had neurites but stained positive for the neural precursor marker Nestin (3.8%, 19 cells of 501 nuclei) or the astrocytic marker glial fibrillary acidic protein (GFAP, 3.7%, 27 cells of 724 nuclei). The enriched glutamatergic identity of these hESC-neurons was substantiated by staining for vesicular glutamate transporter 1 (VGLUT1), a marker of glutamatergic lineage (Fig. 1F). Approximately 38% of all cells were VGLUT1 positive (128 cells of 333 nuclei) and approximately 2.3% were GABAergic, as determined by glutamate decarboxylase (GAD67) immunostaining (12 cells of 516 nuclei). This differentiation efficiency and neuron subtype enrichment are consistent with published hESC-neuron differentiation protocols^25^.

To further evaluate whether the composition of neurites extending into the axonal compartment contained mainly axons, we tested whether a modified rabies virus encoding a fluorescent protein would infect neurons when applied exclusively to the distal neurites within the axonal compartment. Rabies virus infection is specific to axons, requiring endocytosis and retrograde transport of viral particles following attachment to one of three viral receptors located to axons: nicotinic acetylcholine receptor, neuronal cell adhesion molecule or p75 neurotrophin receptor^26^. A modified rabies virus incapable of trans-synaptic transmission and carrying the mCherry gene^27^ was exclusively added to the axonal compartment of hESC-neuron cultures for 2 hours. Once infected through the axon, fluorescent protein expression can be detected throughout the cell, including the axons and dendrites, within 48 hours. Live DIC and fluorescent imaging was performed to visualize the fine processes and fluorescent protein expression throughout the hESC-neurons cultured within the microfluidic chamber (Fig. 1G). While our data suggests these processes were predominantly axons, there is a possibility that dendrites were also were also present in the axonal compartment.

### Differential gene expression between axons and neurons derived from hESCs

On DIV58, after more than 8 weeks in culture, we harvested total RNA from hESC-neurons and isolated neurites within the somatic and axonal compartments of microfluidic chambers (referred to as “neuronal” and “axonal” samples, respectively). The neuronal sample included somata, dendrites and axons. The axonal samples were stringently scrutinized by light microscopy to ensure no somata were present in the axonal compartment. The amount of axonal RNA obtained from individual microfluidic chambers was below detection limits, therefore one round of linear cDNA amplification was performed on a fixed volume of axonal RNA and a fixed mass of neuronal RNA. As a control reaction 10 pg of neuronal RNA was subjected to linear amplification in parallel. The amount of cDNA generated in this reaction suggested that the axonal RNA yield was approximately 2 pg/μL (data not shown). Equivalent amounts of cDNA from all samples was processed for microarray expression profiling (Supplemental Table S1 online), including the 10 pg neuronal sample. Potential chamber-to-chamber variability as well as possible variation in sample collection led us to evaluate the specificity and robustness of our expression profiling. Box-and-whiskers plot of the raw probe cell intensity (log2) by sample shows similarity between the dynamic range of replicates (t-test, p=0.40) (Fig 2A). Pearson’s correlation analysis demonstrated a high degree of correlation between replicates (Fig. 2B). This replicability suggests little variation in the differentiation efficiency between chambers. The 10 pg neuronal sample expression profile more closely resembled the neuronal samples than the axonal samples, suggesting that the linear amplification step broadly preserved the gene expression profile of the low concentration samples.

**Figure 2.**
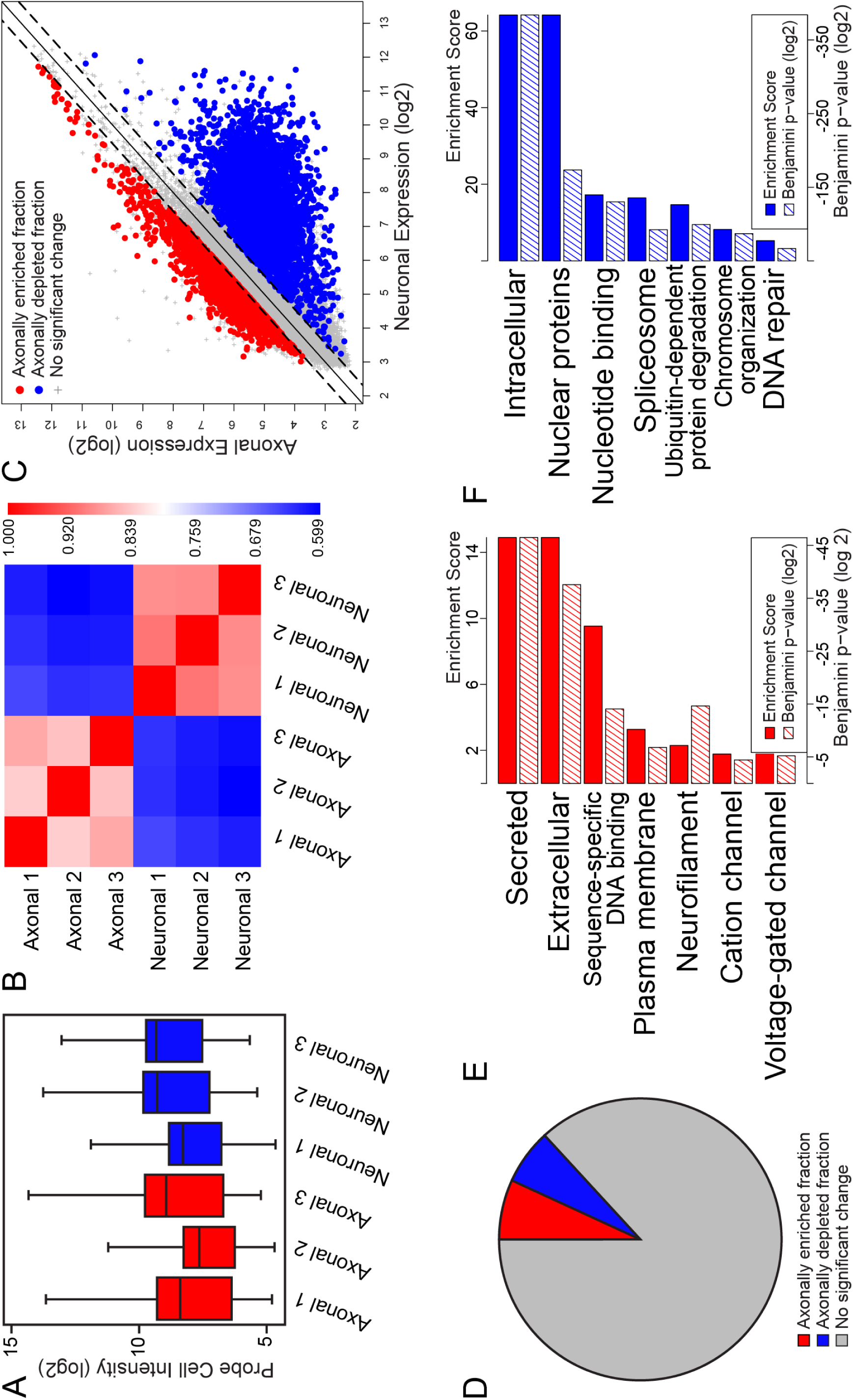
Differential gene expression between the axonal transcriptome and the neuronal transcriptome of hESC-neurons. mRNAs from hESC-neurons grown in axon isolating microfluidic chambers were evaluated using Affymetrix microarrays. (A) Box and whiskers plot of the raw probe cell intensity values (log2) of Affymetrix microarrays of axonal and neuronal samples in triplicate. (B) Pearson correlation coefficient of microarray data from axonal and neuronal samples. (C) Gene expression scatterplot comparing expression levels in hESC-neurons and axons. Threshold for proportional enrichment or depletion within the axonal fraction was set at ±1.5 fold change, ANOVA p-value < 0.05. The proportionally enriched fraction is shown in red, 3942 probe sets. The proportionally depleted fraction is shown in blue, 3630 probe sets. (D) Pie chart depicting the percentages of the total microarray probe sets that reached threshold to be considered enriched (8.17% of probe sets, red region) and depleted (7.53% of probe sets, blue region) within the axonal compartment. (E) Gene ontology (GO) enrichment scores (solid red bars, scale along the top) and Benjamini p-values (hashed red bars, scale along the bottom) of the proportionally enriched axonal fraction as determined by DAVID Gene Functional Classification. (F) GO enrichment scores (solid blue bars, scale along the top) and Benjamini p-values (hashed blue bars, scale along the bottom) of the proportionally depleted axonal fraction as determined by DAVID Gene Functional Classification.

The identification of transcripts within the axonal compartment that have disproportionally high expression may reveal genes that have axonal functions reliant on local translation. We evaluated the proportionally enriched fraction of mRNAs within our axonal dataset by DAVID Gene Functional Classification^28, 29^. For comparison we also evaluated the proportionally depleted mRNAs within the axonal dataset. Figure 2C is a scatterplot of expression levels highlighting these fractions and figure 2D is a pie chart depicting the proportion of probe sets within these two fractions. DAVID Gene Functional Classification of these genes revealed functional categories which were assigned an enrichment score (Fig. 2D, E). This enrichment score measures and weighs the ratio of transcripts that fall into a given functional category. The proportionally enriched mRNAs within hESC-neuron axons were classified as “secreted” and “extracellular” proteins, “DNA sequence-specific binding” proteins, “neurofilament” proteins and “voltage-gated” and “cation” channels (Fig. 2D). This suggests that axonal translation of proteins within these classes may have a more prominent role within the specialized axonal subcellular compartment than within the soma. The proportionally depleted mRNAs were associated with the terms “intracellular” and “nuclear” proteins and proteins that function in RNA splicing, protein degradation and maintaining genome organization and integrity (Fig 2E), functions representative of the identity of this dataset inclusive of somata and dendrites. Taken together these data suggest that enriched transcripts within the axonal samples have the capacity to support a repertoire of functions unique from the cell body.

### Abundant axonally localized transcripts of hESC-neurons functionally resemble axonal transcripts localized to rat cortical neurons

Many of the mRNAs highly abundant within the hESC-neuron axons were not enriched in axons relative to neuronal samples, yet may perform important biological functions that depend on local translation. For example, β-actin (ACTB) is not enriched in axons in our dataset (axonal expression value 6.74, neuronal expression value 6.86, fold change -1.09) nor in the axonal transcriptome of rat cortical neurons, yet local translation of β-actin in axons is well-established ^14, 30^. To assess the degree of functional similarity between the axonally abundant transcripts within hESC-neurons and comparable primary rodent glutamatergic neurons we first created a subset of our axonal expression data containing the microarray probe sets with the highest average axonal signal intensity, those above the 90^th^ percentile (axonal expression value ≥6.48), representing the highest expressed transcripts within the axonal samples (Fig. 3A) (3696 transcripts). *Post hoc* evaluation of this threshold revealed that it excluded known dendritic transcripts, such as ARC, CAMK2A and AMPA receptor subunits and included the well characterized axonal transcripts ACTB (>93^rd^ percentile) and GAP43 (>98^th^ percentile); thus, supporting that our axonal samples contain mostly axons. We compared this subset of highly expressed transcripts by DAVID Gene Functional Classification with a published dataset of mRNAs reliably localized to axons of primary embryonic rat cortical neurons grown in similar axon isolating microfluidic chambers^23^. Transcripts encoding proteins involved in translation and ribosomal proteins were significantly over-represented in the axonally abundant transcripts from both species as well as proteins of the mitochondrial “respiratory chain” and cytoskeletal proteins (“Neurofilament” in hESC-neurons and “Microtubule” and “Cell projection” in rat cortical neurons, Fig. 3C, D). Unique to the hESC-neuron were transcripts encoding “Extracellular” proteins. Overall, these data demonstrate that the axonal transcriptomes of hESC-neurons and primary rat cortical neurons are broadly similar, despite differences in species of origin and neuron derivation.

**Figure 3.**
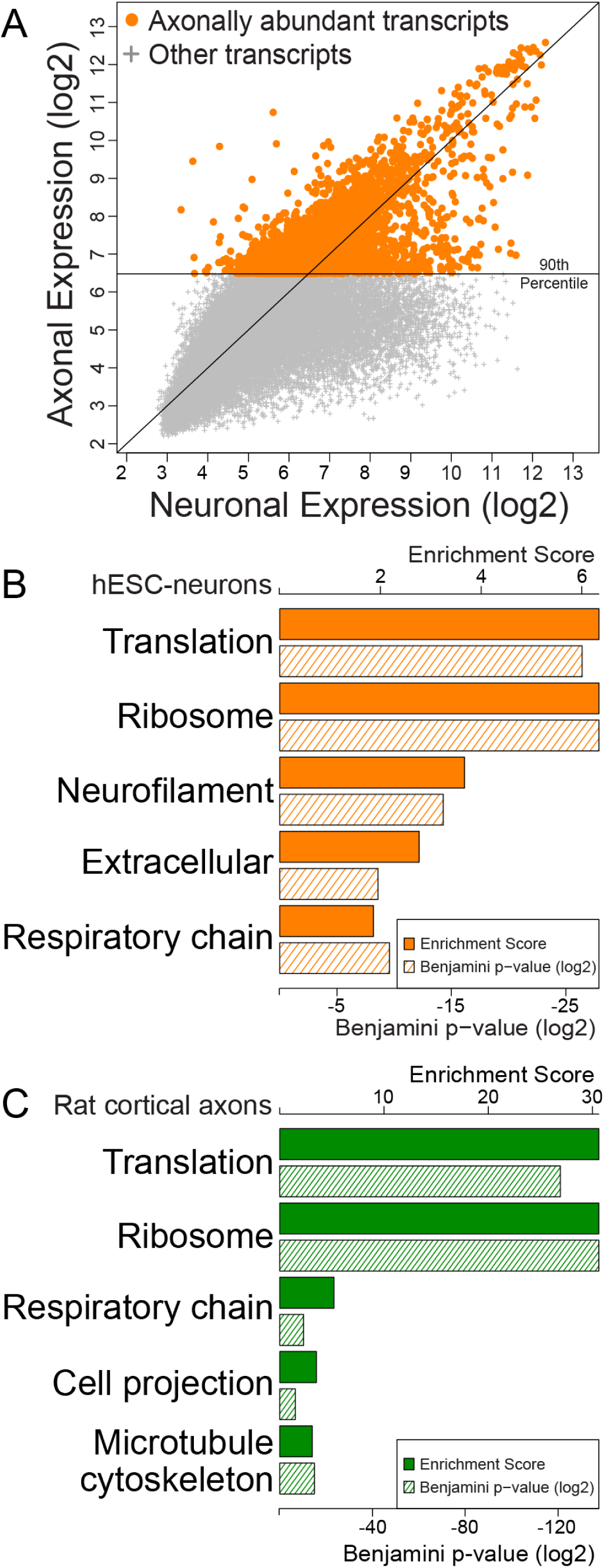
Enriched gene ontology categories were similar between the axonal transcriptomes of hESC-neurons and embryonic rat cortical neurons. (A) hESC-neuron gene expression scatterplot highlighting axonally abundant transcripts with expression levels above the 90^th^ percentile. (B) GO enrichment scores (solid orange bars, scale along the top) and p-values (hashed orange bars, scale along the bottom) of axonally abundant transcripts from hESC-neurons as determined by DAVID Gene Functional Classification. (C) GO enrichment scores (solid green bars, scale along the top) and p-values (hashed green bars, scale along the bottom) of axonal transcripts from rat cortical neurons at 13 DIV^12^ as determined by DAVID Gene Functional Classification.

### RNA-FISH verification of specific mRNAs within hESC-neuron axons

To validate our microarray results, we chose to verify the presence of known axonally localized mRNA transcripts and an unreported mRNA using fluorescence in situ hybridization (FISH). The transcripts selected for visualization were β-actin (ACTB), growth associated protein 43(GAP43) and oxytocin (OXT). ACTB and GAP43 mRNA are well characterized within rodent axons and are locally translated^31^. We identified OXT mRNA as a potentially unique transcript present in these human axons, which is below detection in rodent cortical axons^23^. The presence and function of OXT mRNA within the axon has not been investigated to our knowledge. Due to the short length of the oxytocin mRNA (548nt) and homology with vasopressin mRNA (AVP) our custom OXT RNA-FISH probe set pool (Affymetrix) contained 7 branched DNA sequences instead of the customary 20 sequences, which may explain the smaller size of these fluorescent puncta. Our FISH results confirmed the presence of ACTB, GAP43, and OXT transcripts within hESC-neuron axons (Fig. 4A, C). RNA-FISH probes were omitted for negative control axons (Fig. 4B) and no fluorescence puncta were detected in these samples. Together, these data validate our microarray data and demonstrate that these human axons contain mRNAs also found in axons of model organisms, yet also have mRNAs that may also be uniquely localized to human axons.

**Figure 4.**
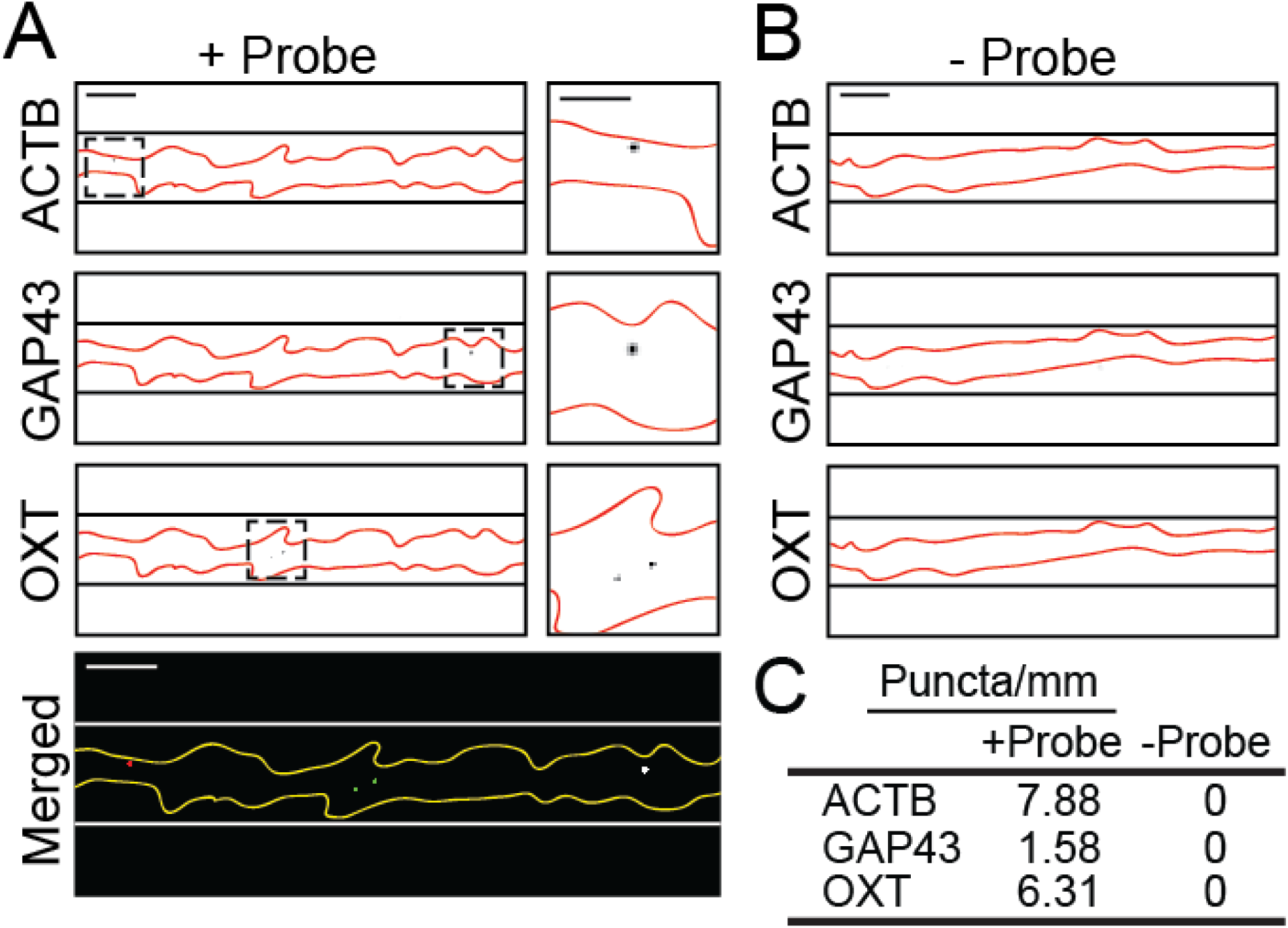
Mulitplexed RNA-FISH verification of mRNA with hESC-neuron axons. (A) Representative portion of hESC-neuron axon processed for RNA-FISH with gene specific probes for β-actin (ACTB, red), Growth Associated Protein 43 (GAP43, green) and Oxytocin (OXT, white). Individual greyscale images (inverted fluorescence) for each probe and a merged color image are shown. In the greyscale images black lines delineate the microgroove walls and the axons are outlined in red. The region within the dashed box is enlarged to the right of each image, scale bar is 5 μm. In the merged image white lines delineate the microgroove walls and the axons are outlined in yellow, scale bars are 10 μm. (B) Representative portion of a negative control hESC-neuron axon processed for RNA-FISH without gene specific probes. Individual greyscale images for each probe are shown. Scale bars 10 μm. (C) Quantification of RNA-FISH puncta per axon length using ImageJ.

## DISCUSSION

Axons, once thought to be devoid of mRNA and ribosomes, are proving to contain transcripts encoding thousands of proteins and the machinery for translation^32, 33^. Functionally, intra-axonal translation is necessary for growth cone guidance, axon maintenance, injury response and may be involved in presynaptic plasticity. Specific locally translated mRNAs involved in these cellular events have been identified^6, 7, 9, 14, 30, 34-43^. Discovery of axonally translated proteins in hESC-neurons has the potential to expand and deepen our knowledge of local translation. While this study focuses on the proportionally enriched fraction and axonally abundant transcripts, many of the previously published axonally translated mRNAs are excluded from these subsets, suggesting intra-axonal translation of moderate to low abundance axonal transcripts may also be functionally relevant.

Functional classification of differentially enriched mRNAs revealed that local translation of retrogradely transported transcription factors (Fig 2C, “Sequence-specific DNA binding”) is likely a common mechanism by which distal axonal events trigger a transcriptional response. Axonal translation of the transcription factors ATF4 and STAT3 are induced following injury in CNS and PNS neurons, respectively^39, 41^. Retrograde transport of these proteins to the soma may mediate the transcriptional response to axon injury. Alternatively, some annotated transcription factors could have transcription-independent roles in the axon. For example, axonally synthesized β-catenin, a Wnt signal transducer, functions at the presynapse as a scaffold protein^9, 39, 41, 44^. The highly enriched category of “secreted” protein transcripts suggests a significant demand on axons to dynamically modulate the extracellular environment, likely as part of axon guidance, synaptogenesis and even synaptic plasticity. Differential expression of the proportionally enriched and proportionally depleted fractions may be the cumulative result of differential trafficking of RNA to the axonal compartment as well as differential mRNA stability within this compartment. Messenger RNA stability can be regulated by multiple mechanisms, including microRNAs and nonsense mediated decay. MicroRNA mediated mRNA regulation is documented in axons^45, 46^. Nonsense mediated decay (NMD) components are present in axons and growth cones and *in vivo* disruption of NMD in mouse commissural neurons altered axon guidance. Spatiotemporal cues developmentally regulate NMD of Robo3.2 mRNA in the axons of these neurons^40^.

Our comparison of the most abundant axonal mRNAs within hESC-neurons with that of primary embryonic rat cortical neurons revealed functional similarities, specifically in mRNAs encoding proteins required for translation, ribosomal proteins and nuclear-encoded mitochondrial proteins necessary for ATP production (Fig. 3B and C). These axon enriched categories have been described in the adult rat neuropil, adult and embryonic rat DRGs, embryonic mouse DRGs, embryonic mouse motoneurons and mouse retinal ganglion cells^10, 24, 47-49^. Axonal translation of nuclear encoded mitochondrial respiratory chain proteins has been described previously^50, 51^. We speculate that it is more efficient to replenish critical mitochondrial proteins via *de novo* and *in situ* translation than for the neuron to support a continual cycle of soma to axon mitochondrial transport. The conservation of ribosomal protein transcripts in axons suggest that either the protein components of the ribosome have a shorter half-life than the RNA components or local translation of ribosomal proteins is a mechanism to dynamically regulate ribosomes and translation. There are over 100 human ribosomal proteins but only a few are constitutive ribosome components and the function of many of these proteins are unknown. It is interesting to speculate that the unique repertoire of ribosomal proteins associated with rRNA could confer target mRNA specificity. Ribosomes containing Ribosomal Protein L38 have been shown to preferentially interact with and translate Hox genes in mouse^52^. With the large number of ribosomal proteins it is possible that this is a common, under characterized, mechanism of translational regulation. Additionally, ribosomal proteins may mediate differential ribosome localization by regulating ribosome interaction with transmembrane proteins of organelles, such as the axonal endoplasmic reticulum, or growth factor receptors, such as the netrin-1 receptor Deleted in Colorectal Cancer (DCC)^53^.

We confirmed the presence of three specific mRNAs, ACTB, GAP43 and OXT, within hESC-neuron axons using multiplexed RNA-FISH. In rodent neurons mRNAs have been shown to be enriched in growth cones but the extremely delicate growth cones of the hESC-neurons were not preserved during the RNA-FISH protocol therefore we were unable to evaluate mRNAs within these structures. It is also possible that the RNA-FISH fixation and protease conditions, which were optimized to preserve the axons, may have been sub-optimal for fixing all mRNAs or unmasking them from ribonucleoprotein complexes. Potentially, but less likely, bias was introduced during RNA sample processing for microarray analysis.

The time course and identifying characteristics of human neuron polarization has not been rigorously investigated so researchers must apply what is known about neuron polarization from other species. Our immunofluorescent evaluation of the neurites within the microgrooves and the axonal compartment demonstrated that they were β-tubulin III positive and MAP2 negative as neurites first extend into the axonal compartment; these are canonical features of properly polarized axons of embryonic and adult neurons from all model species. Further, a modified rabies virus delivered to the isolated axonal compartment was able to infect the hESC-neurons and the GO functional categories were similar between this hESC-neuron axonal transcriptome and the rat cortical axon transcriptome^23^.

Neurons derived from *in vitro* differentiated pluripotent stem cells are generally accepted as resembling fetal neurons^54^. While the differentiation and maturation protocol we used has generated neurons capable of action potentials as early as DIV42^21^, a characteristic of functionally mature neurons, these cells are not “aged” to a point of resembling adult neurons. RNA-mediated mechanisms have been implicated in both neurodevelopmental and neurodegenerative models, and researchers are working to generate appropriately aged stem cell derived neurons^55^.

In the future, the ability to generate patient derived induced pluripotent stem cells and differentiate these cells to neurons in axon isolating microfluidic chambers will facilitate studying axon function in complex genetic diseases such as schizophrenia and autism spectrum disorders. CRISPR-mediated engineered stem cells, such as targeted deletion of RNA binding proteins associated with neurological diseases, applied to our system would provide further insight into the role of axonal translation in human neurons. As the techniques and algorithms necessary to quantitatively evaluate single cell transcriptomes by RNA-seq become robustly established they can be applied to axonal mRNA to evaluate the localization and relative quantities of splice variants, possibly revealing additional levels of complexity within axonal transcriptomes. Further, mechanistic examination of axonal translation in human axons has the potential to reveal relevant axonal mRNAs in disease and normal neuron function.

## METHODS

### Microfluidic chambers

Custom microfluidic chambers were fabricated by soft lithography using an established protocol^22^ with the following modifications. The somatic and axonal compartments of these chambers were 1.5 mm by 7 mm by 450 μm tall. These compartments were connected by microgrooves of 450 μm by 10 μm by 3 μm tall. The only dimension different from previously published microfluidic chambers^23^ was the compartment height, which was increased to facilitate the daily media changes necessary to meet the nutrient demands of hESC-neurons. To create the tall cell compartments a thick layer of photoresist (SU-8-2050; Microchem) was spun on the wafer in 2 coatings. Each coating was spun at 800 rpm for 45 s, the first coating was baked on a leveled 95**°**C hot plate for 3 h and the second coating was baked for 5 h. Wafers were UV-exposed (1000 mJ total over 3 sessions with at least 45 s between exposures) and baked for 1 h in a leveled 95 **°**C oven. Finally wafers were gradually cooled to room temperature over 30 min and developed in PGMEA.

The resulting masters were used to cast microfluidic chambers using poly(dimethylsiloxane) (PDMS) (Sylgard 184 Silicon Elastomer, Dow Corning) as described previously^56^. German glass coverslips were sterilized and coated overnight with a solution of 500-550 kDa poly-D-lysine/laminin (80 μg/ml and 10 μg/ml, respectively), washed and dried. Microfluidic chambers were assembled from PDMS devices and coated coverslips.

### Maintenance and Differentiation of human ESC-neurons

Stem cell differentiation into neural progenitor cells (neuroepithelial cells) was performed according to previously published modifications of the original protocol^20, 21, 57^. Days *in vitro* (DIV) numbering began when H9 human embryonic stem cells were plated in embroid body media to induce neuronal differentiation.

### Maturation of hESC-neurons in microfluidic chambers

On DIV24 mitotic neural progenitor cells were manually lifted and dissociated manually or with Accutase (Life Technologies). Approximately 5x10^3^ cells were seeded into the somatic compartment of microfluidic chambers in N2B27 media supplemented with 100 ng/ml human recombinant brain derived neurotrophic factor (BDNF). Half of the media (100 μl) from each compartment was changed daily and cells were matured for at least 35 days (DIV59). This protocol has been demonstrated to generate glutamatergic neurons as early as DIV25, as verified by staining for the telencephalic transcription factor FOXG1, the transcription factor CTIP specific for subcerebral projection neurons, the glutamatergic transcription factor TBR1 and the glutamatergic marker protein VGLUT1^20, 21^.

### Immunocytochemistry

Human ESC-neurons grown in axon isolating microfluidic chambers were fixed, stained and mounted within the microfluidic chamber because the cells were susceptible to damage when the chambers were removed. Using this method the neuronal processes within the microgroove-embedded barrier region were not exposed to antibodies due to the fluidic isolation of the two compartments. Chambers were fixed with freshly prepared 4% paraformaldehyde in PBS containing 40 mg/ml sucrose, 1 μm MgCl2, and 0.1 μm CaCl2 for 45 min. Neurons were permeabilized in 0.25% Triton X-100 for 15 min then blocked in PBS containing 10% goat serum for 15 min. Primary antibodies to β-tubulin III (alias TUJ1) (1:2000; chicken; Aves Labs), MAP2 (1:1000; rabbit; Millipore), GFAP (1:1000, rabbit; Millipore), Nestin (1:1000, mouse; Abcam) and VGLUT1 (1:200; mouse; NeuroMab) were diluted in PBS with 1% goat serum and incubated overnight at 4 °C. AlexaFluor goat secondary antibodies conjugated to fluorophores with 488 nm, 568 nm, or 633 nm excitation wavelengths (1:1000; Invitrogen) were diluted in PBS and incubated for 1 h at room temperature. Cells were counterstained with DAPI.

### RNA-FISH

Affymetric ViewRNA ISH Cell Assay kit was used for multiplexed visualization of β-actin, oxytocin and GAP43 mRNA in human ESC-neurons grown in axon isolating microfluidic chambers. Commercially available ViewRNA Probe Sets were used to detect human β-actin mRNA (Affymetrix, VA4-10293) and GAP43 mRNA (Affymetrix, VA6-13097) and a custom probe was designed by Affymetrix to detect human oxytocin mRNA (Affymetrix, VA1-20398). The exact same staining procedure was used for the negative control samples, but without the RNA-FISH probes. All steps were performed in a humidified chamber according to manufacturer instructions with the following modifications. Cells were fixed for 30 minutes with freshly prepared 4% paraformaldehyde in PBS containing 40 mg/ml sucrose, 1 μm MgCl2, and 0.1 μm CaCl2. Samples were washed with RNase-free PBS then dehydrated and rehydrated through a series of ethanol washes, as outlined in the ViewRNA ISH Cell Assay optional instructions. The optimal protease treatment was determined to be a dilution of 1:5000 for 3 minutes at room temperature. All subsequent steps followed the Affymetrix ViewRNA ISH Cell Assay user manual.

### Modified rabies virus infection

The axonal compartment of DIV61 cultures were incubated with approximately 100,000 viral units of modified rabies virus carrying the mCherry gene^27^ in a total of 50 μl media for 2 hours at 37 °C, washed twice with fresh media and live imaging was performed 48 hours later. We have observed within 48 hours this viral load results in detectable mCherry expression within 80-85% of hESC-neuron axons in microfluidic chambers. Similar observations were made with primary embryonic rat hippocampal neurons grown in axon isolating microfluidic chambers (data not shown). The presence of mCherry negative axons within the axonal compartment could arise from variability in the time course of mCherry expression. In primary rat hippocampal cultures the minimum amount to time to detect fluorescent protein is 48 hours but some cells do not express detectable mCherry until 4 days after transfection.

### Confocal imaging

Z-stack fluorescent images and single plane differential contrast (DIC) images were acquired using a spinning disk confocal imaging system (Revolution XD, Andor Technology) configured for an Olympus IX81 zero-drift microscope and spinning disk unit (CSU-X1, Yokogawa). Light excitation was provided by 50 mW, 488 nm; 50 mW, 561 nm; and 100 mW, 640 nm lasers. The following bandpass emission filters (BrightLine, Semrock) were used: 525/30 nm (TR-F525-030), 607/36 nm (TR-F607-036), 685/40 nm (TR-F685-040). All slides within a series were imaged during a single session in which image capture settings were identical for all fields.

### Image analysis

Z-stack fluorescent images were MAX projected. 16-bit images were manually thresholded to the same values within each series of stained chambers.

### Determination of differentiation efficiency

Seven fields of hESC-neurons in monolayer from 4 independently stained chambers of at least DIV39 were used to estimate the efficiency of differentiation to mature hESC-neurons as detected by colocalization of DAPI signal with MAP2 or β-tubulin III positive staining. Three fields from two independently stained chambers of at least DIV49 were used to evaluate Nestin, GFAP, VGLUT1 and GAD67.

### RNA isolation

On DIV59 isolation of RNA from microfluidic cultures using the RNAqueous-Micro Kit (Ambion) was performed as previously described^22, 23^. To collect pure axonal RNA continuous aspiration was applied to the somatic compartment to prevent somatodendritic RNA from entering the sample. Most of the axonal media was removed and lysis buffer was added to one well, the reagent that flowed to the other axonal well through the axonal compartment was collected as the axonal RNA sample. Subsequent RNA purification was performed according to manufacturer’s instructions and RNA was eluted in a final volume of 10 μl. Neuronal RNA was collected from the somatic compartment of hESC-neurons grown in microfluidic chambers not used for axonal RNA collection, this compartment contained somata, dendrites and axons which did not pass through the microgrooves. The majority of the media from the somatic compartment was removed and the same RNA isolation procedure was followed, omitting constant aspiration. Each sample was obtained from one microfluidic chamber. Samples were stored at -80°C until further processing by the UNC Lineberger Comprehensive Cancer Center Genomics Core and the UNC School of Medicine Functional Genomics Core. Three axonal and three neuronal samples were collected and analyzed.

### RNA amplification and microarray

Axonal RNA harvested from microfluidic chambers is below the detection limit of currently available technology. A fixed volume of each axonal RNA sample (5 μL) and a fixed mass of each neuronal RNA sample (250 pg) were subjected to one round of linear amplification using the Ovation One-Direct System (NuGEN). This system initiates amplification at the 3’ end as well as randomly throughout the transcript and is optimized for very small biological samples such as single cell transcriptomics. The integrity and concentration of the resulting cDNA samples were quantified with an Agilent Bioanalyzer 2100 at the UNC Lineberger Comprehensive Cancer Center Genomics Core. After amplification axonal samples were 230.5 ng/μl, 305.0 ng/μl and 199.0 ng/μl. Neuronal samples were 519.7 ng/μl, 497.3 ng/μl and 469.9 ng/μl. As a control reaction, 10 pg of neuronal RNA was amplified resulting in a concentration of 252.1 ng/μl, suggesting our original axonal RNA yield ranged from approximately 1.6 to 2.4 pg/μl. A fixed concentration of amplified cDNA from all samples, including the 10 pg neuronal sample, were processed in parallel for microarray analysis using the Encore Biotin Module (NuGEN) at the UNC School of Medicine Functional Genomics Core.

Human Gene 2.0 ST Arrays (Affymetrix) were used and scanned with an Affymetrix GeneChip Scanner 3000 7G Plus with Autoloader. Affymetrix Expression Console software was used to analyze microarray CEL files. Affymetrix normalization controls, built into the microarray chips and analysis software, were used for raw data normalization. The Robust Multichip Analysis (RMA) algorithm was used for global background adjustment, quantile normalization and gene expression summarization between samples. This method is sensitive to small changes between samples without negatively affecting signal variance. Affymetrix Transcriptome Analysis Console v2.0 software algorithms were used to determine differential expression and statistical analysis using one-way between-subject ANOVA of normalized intensities (Supplemental Table S1 online).

### Data analysis

Microarray probe sets with fold change linear (Axon vs Neuron) greater than or equal to 1.5 and an ANOVA p-value (Axon vs Neuron) less than 0.05 were designated the axonally enriched fraction (3943 probe sets). Probe sets with fold change linear (Axon vs Neuron) less than or equal to -1.5 and an ANOVA p-value (Axon vs Neuron) less than 0.05 were designated the axonally depleted fraction (3631 probe sets). The fold change threshold of 1.5 was selected to counter balance the compressed calculated fold change values obtained by the RMA method. This threshold generated gene symbol lists of an appropriate size for DAVID Functional Annotation Clustering (<3000). Official gene symbols were extracted, duplicates were removed and gene symbol lists (3297 for axonally enriched fraction and 3638 for axonally depleted fraction, Supplemental Tables S2 and S3 online, respectively) were submitted to DAVID Bioinformatics Resources for Gene ID Conversion and Functional Annotation Clustering with the single species *Homo sapiens* selected and the default human background list^28, 29^.

To obtain a gene symbol list of axonally abundant transcripts within hESC-neuron axons microarray probe sets with Axon bi-weight average signal (log2) equal to or greater than 6.48 were selected. This threshold excluded known dendritic transcripts and included β-actin, a well characterized axonal mRNA. These criteria were similar to those used by Taylor *et al.* (2009) to determine axonal mRNAs within primary rat cortical neurons. Official gene symbols were extracted, duplicates were removed and the gene symbol list (3696 gene symbols, Supplemental Table S4 online) was submitted to DAVID Bioinformatics Resources for Gene ID Conversion and Functional Annotation Clustering with the single species *Homo sapiens* selected and the default human background list^28, 29^.

This list of 3696 human gene symbols was compared to published gene symbols of axonally localized transcripts from rat cortical neurons as obtained through the NCBI Gene Expression Omnibus, GEO number GSE11730, and in published supplementary data^23^.

## ACKNOWLEDGEMENTS

The authors would like to acknowledge the CHANL facility at UNC-CH. R.L.B. was supported in part by a grant from the National Institute of General Medical Sciences under award 5T32 GM007092. A.M.T. acknowledges support from the NIH (R42MH097377) and the Simons Foundation (SFARI #236390).

## AUTHOR CONTRIBUTIONS

R.L.B., J.W.K., R.D., M.N. and A.M.T. designed and performed experiments. R.L.B. and A.M.T. prepared the figures and wrote the manuscript.

## ADDITIONAL INFORMATION

### Accession codes

All microarray data have been deposited to the NCBI Gene Expression Omnibus under accession number GSE84975, NCBI tracking system ID 17986454.

